# Blueberry and cranberry pangenomes as a resource for future genetic studies and breeding efforts

**DOI:** 10.1101/2023.07.31.551392

**Authors:** Alan E. Yocca, Adrian Platts, Elizabeth Alger, Scott Teresi, Molla F. Mengist, Juliana Benevenuto, Luis Felipe V. Ferrão, MacKenzie Jacobs, Michal Babinski, Maria Magallanes-Lundback, Philipp Bayer, Agnieszka Golicz, Jodi L Humann, Dorrie Main, Richard V. Espley, David Chagné, Nick W. Albert, Sara Montanari, Nicholi Vorsa, James Polashock, Luis Díaz-Garcia, Juan Zalapa, Nahla V. Bassil, Patricio R. Munoz, Massimo Iorizzo, Patrick P. Edger

**Author notes:** Luis Felipe V. Ferrão. David Chagné. Luis DÍaz l.

## Abstract

Domestication of cranberry and blueberry began in the United States in the early 1800s and 1900s, respectively, and in part owing to their flavors and health-promoting benefits are now cultivated and consumed worldwide. The industry continues to face a wide variety of production challenges (e.g. disease pressures) as well as a demand for higher-yielding cultivars with improved fruit quality characteristics. Unfortunately, molecular tools to help guide breeding efforts for these species have been relatively limited compared with those for other high-value crops. Here, we describe the construction and analysis of the first pangenome for both blueberry and cranberry. Our analysis of these pangenomes revealed both crops exhibit great genetic diversity, including the presence-absence variation of 48.4% genes in highbush blueberry and 47.0% genes in cranberry. Auxiliary genes, those not shared by all cultivars, are significantly enriched with molecular functions associated with disease resistance and the biosynthesis of specialized metabolites, including compounds previously associated with improving fruit quality traits. The discovery of thousands of genes, not present in the previous reference genomes for blueberry and cranberry, will serve as the basis of future research and as potential targets for future breeding efforts. The pangenome, as a multiple-sequence alignment, as well as individual annotated genomes, are publicly available for analysis on the Genome Database for Vaccinium - a curated and integrated web-based relational database. Lastly, the core-gene predictions from the pangenomes will serve useful to develop a community genotyping platform to guide future molecular breeding efforts across the family.

## Introduction

The heath family (Ericaceae) contains many culturally and economically important berry crops, including bilberry (*Vaccinium myrtillus* L.), blueberry (*Vaccinium spp*. L.), cranberry (*Vaccinium macrocarpon* Aiton), huckleberry (*Vaccinium membranaceum* Douglas ex Torr.) and lingonberry (*Vaccinium vitis-idaea* L.) ^1^. The common name ‘blueberry’ is applied to multiple *Vaccinium* species, including highbush blueberry (*V. corymbosum* L.), lowbush blueberry (*V. angustifolium* Aiton), and rabbiteye blueberry (*V. virgatum* Aiton [synonym = *V. ashei* J.M. Reade]) ^2,3^. Worldwide demand and consumption of cranberries and blueberries has rapidly increased over the past decades in large part owing to their health-promoting properties ^4,5^. Both cranberry and highbush blueberry are native to North America ^6,7^. Cranberry is a diploid species (2n=2x=24) whereas highbush blueberry is a tetraploid (2n=4x=48) ^8,9^. Highbush blueberry is further subdivided into northern and southern varieties that were selected largely for their overall chilling requirements for flowering and winter hardiness differences ^10^. The cultivation of cranberry and highbush blueberry began in the early 1800s and 1900s, respectively ^11,12^. Thus, the domestication history of these species is much shorter than those of other crops and there remains a great unexplored breeding potential for all *Vaccinium* crops ^13^.

Recent studies have shown that sequencing a single reference genotype or cultivar of a species is insufficient to recover all the genetic diversity present in a group ^14–17^. This was first recognized in microbial studies; sets of genes were found to either be present in every member of a population (core) or absent in at least a single individual (dispensable) ^18^. Here, we choose to refer to dispensable genes as auxiliary genes ^19^. Although absent in some individuals, certain auxiliary genes if lost in combination can be lethal because of either redundancy or epistatic interactions with other auxiliary genes ^20^. The sum of all core and auxiliary genes is termed a pangenome ^21^. Several pangenome studies have been conducted in plants including *Brachypodium distachyon, Brassica napus*, maize, soybean, rice, and strawberry ^22–27^.

Core genes are consistently found to be enriched for “housekeeping” functions including essential metabolic processes, whereas auxiliary genes were found to be enriched for more adaptive functions. For example, in *Brassica oleracea* (European cabbage), auxiliary genes are strongly enriched for defense response and specialized metabolism, including those that contribute to unique flavor profiles and vitamin content ^28^. In *Brachypodium distachyon*, auxiliary genes were similarly enriched for defense functions as well as other adaptive traits that are of particular interest for breeding superior crop varieties ^22^. In addition, auxiliary genes often display signatures of elevated sequence turnover and relaxed selection and are shorter relative to core genes ^29^.

The proportion of core and auxiliary genes identified can vary for a given species or crop type. This value ranges from 33% to 80% of core genes. For example, about 80% of genes in rice (*Oryza sativa*) are core, while in corn (*Zea mays*), roughly 40% of genes are core ^27,30^. For a summary of core genes predictions across several plant pangenome studies see Golicz et al, 2020 ^17^. Life history characteristics and representative divergence likely contribute to the relative amount of auxiliary genes present for a particular species pangenome ^31^. Furthermore, the rates of structural variation also contribute to pangenome size characteristics ^32^.

Characterization of auxiliary genes is crucial to maximize the impacts of molecular breeding approaches ^33^. Genes underlying many important target traits (e.g. metabolites associated with fruit quality or disease resistance) are often auxiliary ^24^. Previous genome-wide association studies (GWAS) have uncovered several additional candidate loci controlling traits of interest when leveraging a pangenome ^31,34^. For example, one GWAS in pigeon pea uncovered a gene associated with seed weight that was absent in the primary reference genome ^35^. Similarly, Song et al. (2020) leveraged presence-absence variation information from eight reference quality *Brassica* genomes to perform a GWAS and identified novel transposable element insertions associated with variation in flowering time and silique weight ^36^. This illustrates the translational impact of extending analyses beyond single reference genome frameworks.

Here, we generated and annotated genomes for ten diverse cranberry and twenty diverse highbush blueberry cultivars. In conjunction with a previously published reference genome for these crops ^37–39^, we developed a pangenome for these crop species separately, and combined for both species and estimated the core genome size across the genus *Vaccinium*. In addition, we explored distinguishing features between core and auxiliary genes in blueberry and cranberry. These genomic resources and our pangenome estimates will serve as a powerful resource to guide future molecular breeding efforts and genetic studies across the *Vaccinium* community.

## Results

### Selection of accessions, sequencing, assembly, and annotation

We selected ten cranberry cultivars, ten southern highbush blueberry cultivars, and ten northern highbush blueberry cultivars for genome sequencing and annotation. For cranberry and blueberry, reference genomes for cultivars ‘Stevens’ and ‘Draper’, respectively, were published previously and included in our analyses ^38,40^. Cultivars were selected based on genetic marker and pedigree analysis to capture the greatest amount of diversity ^41^ (Figure S1). Accessions were sequenced to an average depth of 112.5X for cranberry and 53.8X for blueberry across each haplotype or 225X for cranberry and 215.2X for blueberry in comparison to a single haplotype reference genome (Table S1). A hybrid reference-based and *de novo* assembly method was used to assemble genomes for each individual^42^. To guide annotation, RNA-seq data were collected from leaf and berry tissue. Each of the genomes were annotated using MAKER2, using the aforementioned RNAseq data, producing on average 27,856 and 105,523 genes for cranberry and blueberry, respectively. Our genome assembly qualities closely reflect those of the ‘Stevens’ and ‘Draper’ reference genomes ^38,40^. ‘Draper’ is a haplotype phased reference genomes, which is thus why roughly four times as many genomes were identified for blueberry than for cranberry. Scaffold N50 values were ∼37Mb for northern highbush genomes, ∼1.4Mb for southern highbush genomes, and ∼38Mb for cranberry genomes (Table S1). Complete BUSCO scores ranged from 82.9% to 91.7% (Table S1).

### Identification of cranberry core and auxiliary genes

All eleven cranberry genomes were aligned using ProgressiveCactus^43^. To identify core and auxiliary genes in cranberry, we integrated results from Orthofinder2 and our genome alignment^44^. For core gene classification, we search for genes present in each individual. Orthofinder2 alone will identify the presence of gene family members. Therefore, to integrate syntenic information, we filtered for the presence of alignments in our ProgressiveCactus results. Since annotations and genomes were generated for all eleven cranberry genotypes, we could label every gene (302,090 total) as either core or auxiliary. Of the roughly 27,463 genes, on average per accession, 14,553 (53%) and 12,910 (47%) genes were identified as core and auxiliary respectively (**Figure 1**).

**Figure 1:**
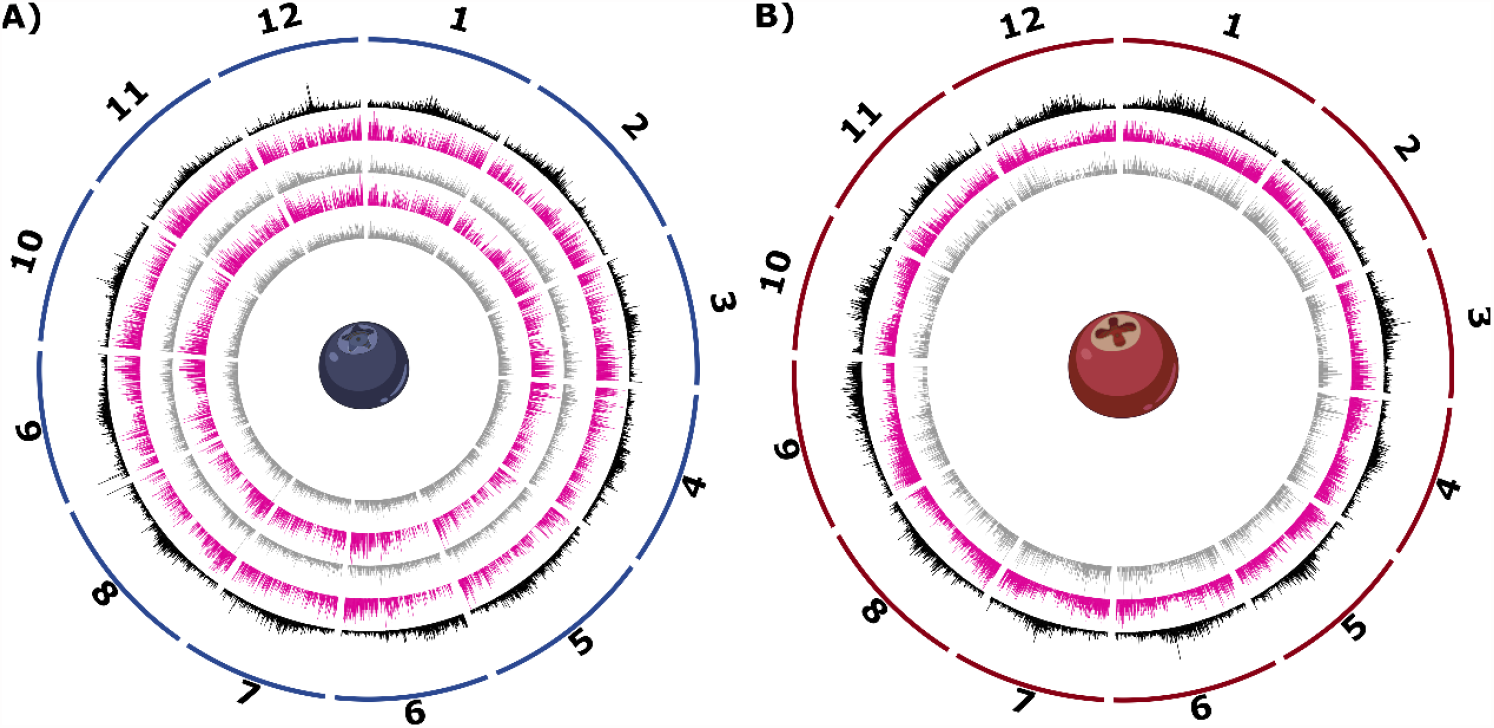
Circos plots for the blueberry and cranberry pangenomes. Circos plots for the blueberry (panel A) and cranberry (panel B) pangenomes are shown. Tracks from exterior to interior are as follows: (1) karyotype of the 12 main pseudomolecules, (2) density of LTR transposons in black, (3) density of core genes in pink, and (4) density of auxiliary genes in gray. For panel A, northern highbush blueberry is shown on the third and fourth tracks, while the two innermost tracks reflect southern highbush blueberry core and auxiliary genes.

### Identification of blueberry core and auxiliary genes

Highbush blueberry is an autotetraploid and the ‘Draper’ genome assembly for highbush blueberry consists of four haplotypes ^37^. Therefore, calling core and auxiliary genes is complicated by the differential presence or absence of genes on different haplotypes. Gene fractionation (loss) in polyploid genomes is expected and will inflate auxiliary designations ^45^. To facilitate core and auxiliary gene designations, we compared our blueberry assemblies with a recently published diploid genotype (W85) for blueberry ^39^ to accurately assess the presence or absence of each gene in the northern and southern highbush blueberry genome assemblies. The W85 genome assembly consists of phased haplotypes with 34,848 gene models in the primary haplotype (p0) and 33,148 in the alternate haplotype (p1). As pangenome patterns were largely consistent between haplotypes, the rest of our analyses focused on the primary “p0” haplotype. Of these, 45.80% (n = 15,150) were present in all highbush blueberry cultivars based on purely genome synteny analyses. Northern highbush blueberry exhibited a higher proportion of core genes (53.87%) than southern highbush blueberry (48.78%), and similar numbers were estimated for each haplotype. We also analyzed core genome size using OrthoFinder2, as we did above with cranberry, and identified 14,956 (51.6%) as core orthogroups and 14,042 (48.4%) auxiliary orthogroups shared across all highbush blueberry cultivars (**Figure 2**). Orthogrouping permitted us to identify core genes that were no longer in the ancestral position but still present in the genome.

**Figure 2:**
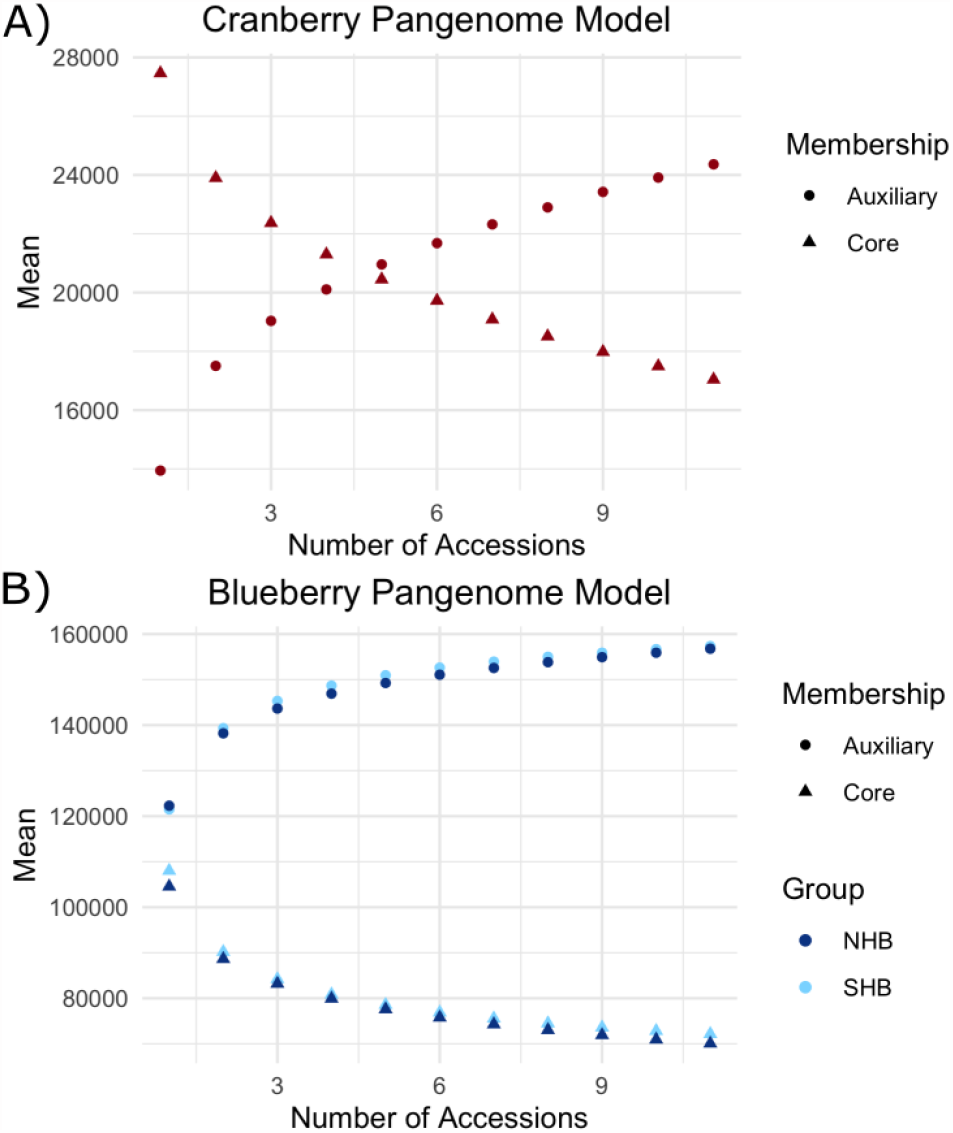
Core and auxiliary genome modeling for cranberry and blueberry. Panel A depicts the model for the core (circle) and auxiliary (triangle) pangenome for cranberry. For each point along the x-axis, we take every possible combination of that size from our genome samples and plot the average number of core and auxiliary genes as a point. Panel B depicts the model for the core (triangles) and auxiliary genes (circles) for northern (dark blue) and southern (light blue) highbush blueberry. Note the y-axes do not start at zero.

### Pangenome modeling

Though cultivars were selected to capture the greatest genetic diversity (**Figure S1**), to capture all genetic diversity we must assay every individual. However, we can estimate the size of the pangenome through modeling our sample. **Figure 2** displays this model of the core and auxiliary gene content of the pangenome estimated using orthogrouping to best capture core and auxiliary genes that were no longer in their ancestral position in the genome. For cranberry, we identified 14,552 core genes out of an average of 27,462 total genes per accession (53%). The number of total auxiliary genes increases as more genomes are queried. However, the amount of new auxiliary genes per accession added decreases. The pangenome is considered “closed” when the total number of auxiliary genes eventually reaches a plateau. That plateau has not been reached modeling on our data. Therefore, further sampling of cranberry genomes will uncover greater genetic variation and additional novel auxiliary genes.

For blueberry, the model of auxiliary and core gene content appeared to reach a plateau more quickly than cranberry. This suggests the same numbers of blueberry individuals for both northern and southern highbush varieties captures a greater proportion of total auxiliary genes than for cranberry, and that we are closer to estimating the true core genome size for blueberry. However, we still observe an incomplete plateau and expect to uncover novel auxiliary genes and an improved core genome size as more genomes are queried. In northern highbush blueberry, there were an average of 70,073 core genes out of 104,552 total genes per accession (67.02%). In southern highbush blueberry, there were an average of 72,155 core genes out of an average of 108,020 total genes per accession (66.80%).

### Differences between auxiliary and core genes

Previous studies identified differences between core and auxiliary genes including gene length, exon count, and guanine-cytosine (GC) percentage ^29,46^. Here, we uncovered similar differences between these two groups of genes in both cranberry and blueberry. **Figure 4** displays differences between core and auxiliary genes across cranberry (CB), northern highbush blueberry (NHB) and southern highbush blueberry (SHB). Auxiliary genes are dramatically shorter, and have both fewer and shorter introns than core genes. Furthermore, we found that the expression of core genes was higher on average than that of auxiliary genes (Figure 3).

**Figure 3:**
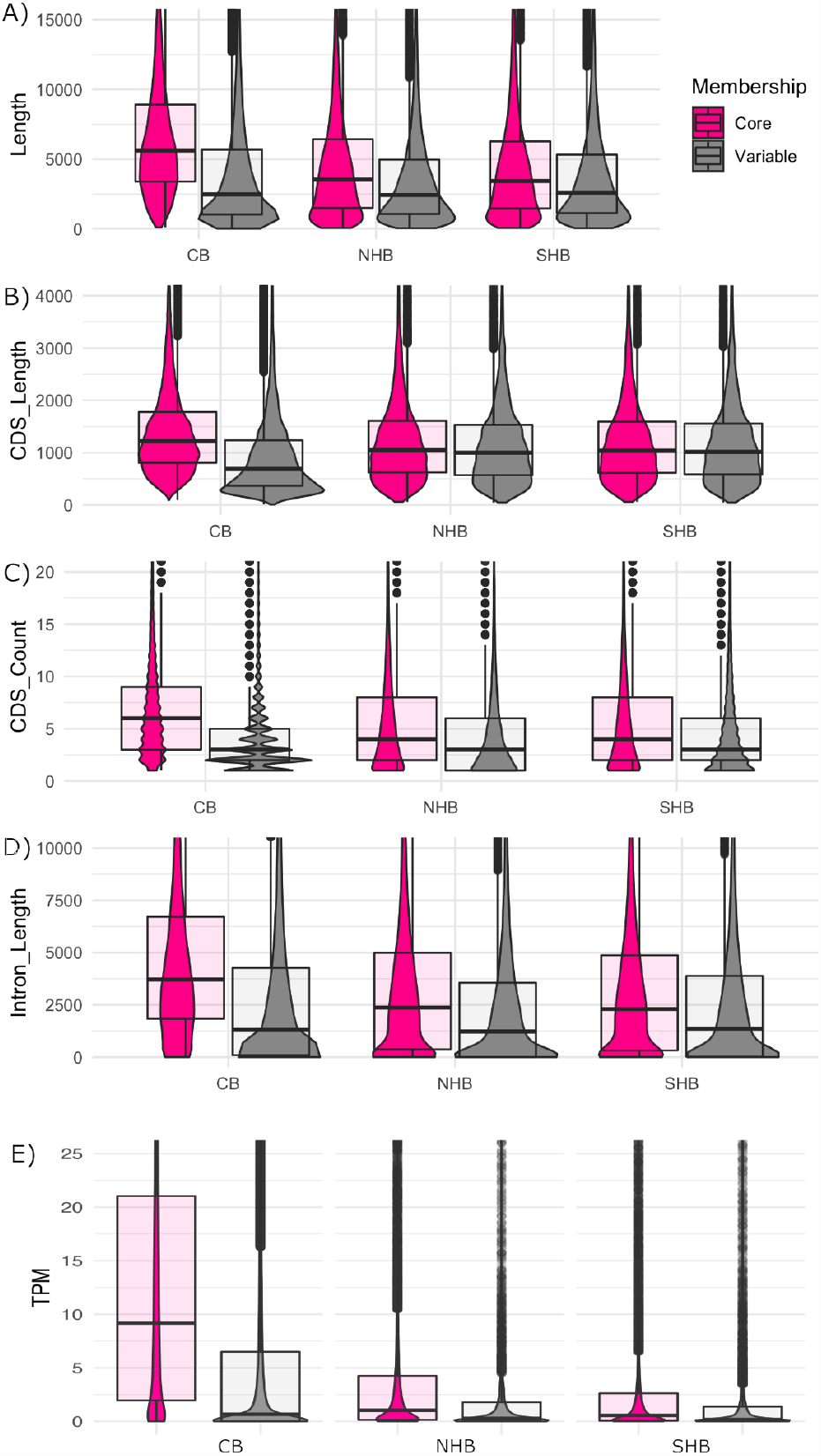
Differences between core and auxiliary genes. Density plots showing the differences between core (pink) and auxiliary (gray) genes for cranberry (CB), northern highbush blueberry (NHB) and southern highbush blueberry (SHB). We compared (A) gene length (transcription start site to transcription end site), (B) coding sequence length (CDS) length, (C) CDS count, (D) intron length, and (E) fruit transcripts per million (TPM).

**Figure 4:**
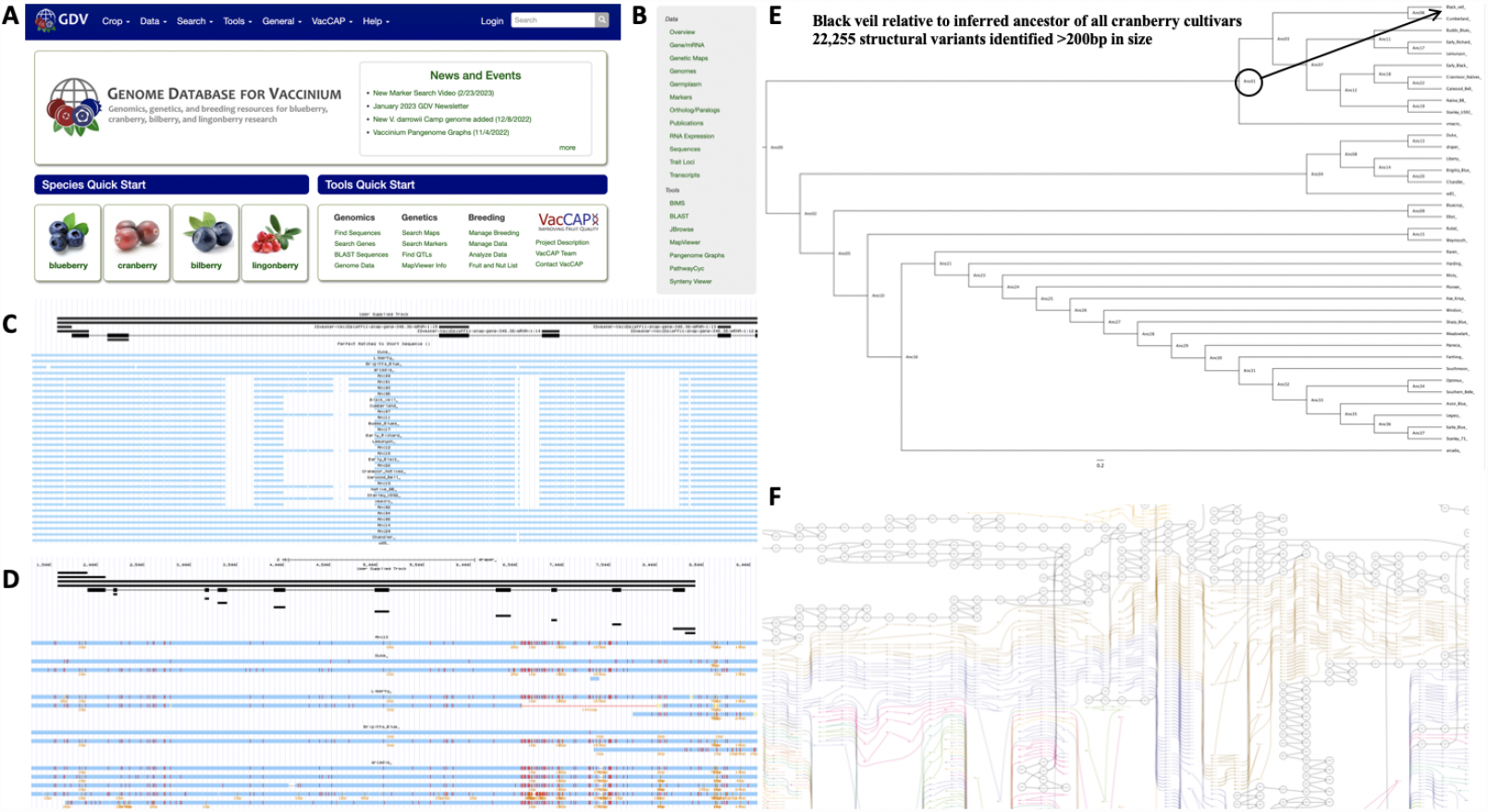
Pangenomic resources available on Genome Database for *Vaccinium*. All genomic resources developed here are publicly available for the community to use on the Genome Database for Vaccinium (GDV; https://www.vaccinium.org/) (Panel A). A wide variety of tools are available to analyze individual genomes and/or compare genomes (Panel B). A multiple sequence alignment is available to identify structural variants, including deletions (Panel C), single nucleotide polymorphisms (Panel D), and place genetic variants into a phylogenetic context (Panel E). Panel F depicts a pangenome graph view of the multiple sequence alignment.

Next, we compared functional differences between core and auxiliary genes (Table S2). For auxiliary genes in cranberry, we did observe a few expected enriched GO terms that were reported in previous studies ^42^ with the top three GO terms being “GO:0009607 response to biotic stimulus” (FDR p-value = 1.22E-08), “GO:0043207 response to external biotic stimulus” (FDR p-value = 1.66E-08), and “GO:0051707 response to other organism” (FDR p-value = 1.66E-08). In addition, auxiliary genes were enriched with many other GO terms of significance to important target breeding traits, including “GO:0009631 cold acclimation” (FDR p-value = 0.0353), “GO:0002213 defense response to insect” (FDR p-value = 0.0041), and “GO:0050832 defense response to fungus” (FDR p-value = 0.0031). Furthermore, we observed enrichment for several GO terms relevant to flavonoids, a group of specialized metabolites, previously shown to affect the pigmentation, health benefits (antioxidant), and defense against pathogens in berries ^47^ (e.g. “GO:0009812 flavonoid metabolic process” (FDR p-value = 0.0388), “GO:0051555 flavonol biosynthetic process” (FDR p-value = 0.0023)). For cranberry, core genes were enriched for core biological processes, with the top three enriched GO terms being “GO:0019222 regulation of metabolic process” (FDR p-value = 3.37E-08), “GO:00104 regulation of gene expression” (FDR p-value = 8.36E-07), and “GO:0065007 biological regulation” (FDR p-value = 1.76E-06).

Similarly, in blueberry, the top three enriched GO terms for auxiliary genes were “GO:0006952 defense response” (FDR p-value = 6.10E-17), “GO:0009607 response to biotic stimulus” (FDR p-value = 1.03E-14), and “GO:0044419 biological process involved in interspecies interaction between organisms” (FDR p-value = 1.04E-14). In addition, auxiliary genes were enriched with many other GO terms of significance to important target breeding traits, including “GO:0071497 cellular response to freezing” (FDR p-value = 0.0279), “GO:0050832 defense response to fungus” (FDR p-value = 6.61E-07), and “GO:0009617 response to bacterium” (FDR p-value = 4.49E-12). Furthermore, several GO terms relevant to the pigmentation, health benefits, aroma and flavor of berries are enriched among auxiliary genes (e.g. “GO:0009813 flavonoid biosynthetic process” (FDR p-value = 0.0009), “GO:0019745 pentacyclic triterpenoid biosynthetic process” (FDR p-value = 0.0249), and “GO:0042335 cuticle development” (FDR p-value = 0.0037)). For core genes in blueberry, the top three enriched GO terms were “GO:0006396 RNA Processing” (FDR p-value = 3.76E-31), “GO:0034641 cellular nitrogen compound metabolic process” (FDR p-value = 1.26E-21), and “GO:0034660 ncRNA metabolic process” (FDR p-value = 4.78E-21).

### Individual genomes and pangenome resources available on public database

Each of the assembled highbush blueberry and cranberry genomes, alongside gene and repeat annotations, are now publicly available on the Genome Database for Vaccinium ^48^ (**Figure 4A**). The Genome Database for Vaccinium (GDV) is a curated and integrated web-based relational database to house and integrate genomic, genetic and breeding data for *Vaccinium* species. Members of the community can visit GDV to view a gene(s) of any genome in JBrowse, search for one or more sequences using BLAST, use synteny viewer to display all the conserved syntenic blocks between any genomes, and pathway predictions including for those that encode specialized metabolites associated with superior fruit quality (**Figure 4B**). For example, presence-absence variation of genes involved in the biosynthesis of flavonoids was recently assessed across the blueberry pangenome ^49^. In addition, the multiple whole genome alignment is available that can be used to view structural variants, including insertion/deletions (InDels)(**Figure 4C**), and small polymorphisms (**Figure 4D**). ProgressiveCactus ^43^ was also used to estimate ancestral states along a phylogeny of the sequenced genome, which can be leveraged to determine the number of events that occurred along a particular branch. For example, we identified 22,255 structural variants >200bp that occurred since the most common recent ancestor of all eleven cranberry and present in ‘Black Veil’ (**Figure 4E**). A complete list of all identified variants, ranging from small single nucleotide substitutions to larger structural events (insertions, deletions, inversions, duplications, and transpositions) are available in Table S3. In addition, pangenome variation graphs for blueberry and cranberry loci can be generated from the hierarchical alignment (HAL) on the GDV (see for example **Figure 4F**).

### Pangenome conservation and as a new resource for the development of future breeding tools

Blueberry and cranberry species belong to two distinct and distantly related clades that diverged between 5 and 10 million years ago (**Figure 5A**) ^50^. We evaluated the conservation of positionally conserved core genes between highbush blueberry and cranberry, which revealed a Blueberry-Cranberry “BC Core” consisting of 10,230 genes (**Figure 5B**). A total of 4,920 and 4,323 core genes were identified as highbush blueberry and cranberry specific, respectively. Next, we evaluated the conservation of the “BC Core” gene content in comparison to other *Vaccinium* species with available genomes. In the small cranberry (*V. microcarpum*) genome ^38^, roughly 95.5% (9,767 total) genes from the “BC Core” were identified and positionally conserved in the genome. For bilberry (*V. myrtillus*) ^51^ and Darrow’s blueberry (*V. darrowii*) ^52^, roughly 94.5% (9,672 total) and 91.3% (9,345 total) genes, respectively, from the “BC Core” were identified and positionally conserved in the genomes. A major goal of the VacCAP project ^53^, a multi-institutional and multi-disciplinary project, is the assembly of a pangenome in order to construct a robust genotyping platform that can be used for molecular breeding efforts for blueberry, cranberry and potentially other related cultivated *Vaccinium* species. Here, we identified a set of 10,230 positionally conserved core genes that span each chromosome of both blueberry and cranberry species, and are largely present (>90%) in related *Vaccinium* species in positionally conserved regions of the genome.

**Figure 5:**
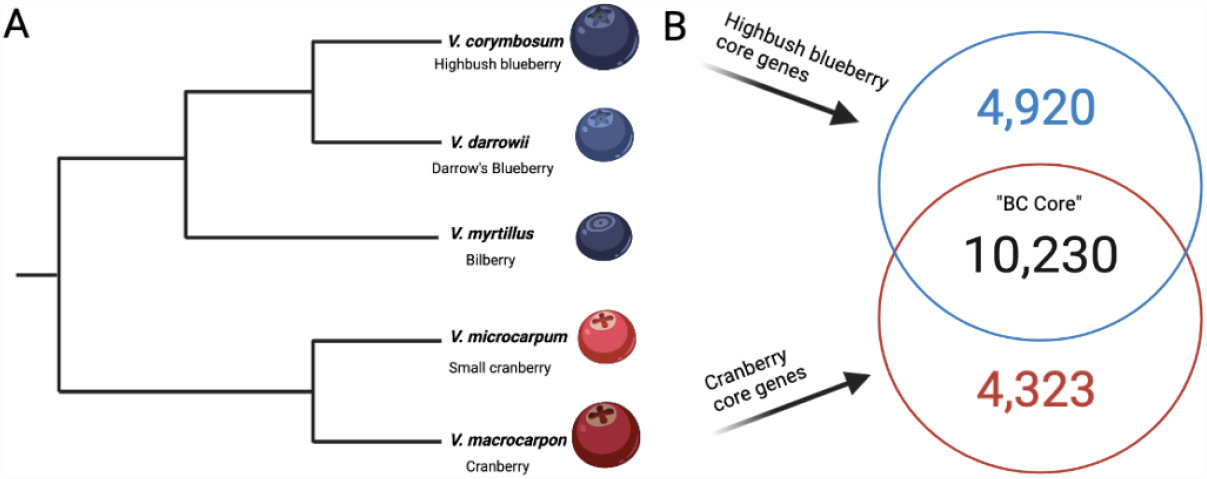
Conservation of highbush blueberry-cranberry (BC) core gene content. Panel A depicts the phylogenetic relationships among five *Vaccinium* species, previously estimated by ^50^. Panel B depicts the comparison of the highbush blueberry and cranberry syntenically conserved core genes.

## Discussion

In this study, we defined genes in highbush blueberry and cranberry as core or auxiliary based on their presence or absence across diverse cultivars. In cranberry, we uncovered 53% of all genes being core and 47% as auxiliary genes. In blueberry, we uncovered similar rates, with 51% of all genes being core and 49% as auxiliary genes. The proportions of core and auxiliary genes identified for blueberry and cranberry were similar to those of other crops and species, including maize ^27^, *Brachypodium* ^22^ and strawberry ^26^. However, not all crop species exhibit presence-absence variation that impacts roughly half of the gene content. Three main factors affect the proportion of core genes that may be identified for a species: divergence time of the genotypes compared, extant diversity of the species and life history traits.

Previous studies have also identified characteristic differences between core and auxiliary genes ^29^. These differences lend insight into core/auxiliary gene function as well as their origin and subsequent evolution. Both core and auxiliary gene characteristic distributions overlap. Therefore, dichotomies cannot be drawn between these two classes. Rather, as with most biological processes, they exist on a continuum. That being said, our identified characteristic differences tell us auxiliary genes show patterns of evolutionarily young genes and are enriched with more adaptive functions that are important breeding targets.

Shorter sequences for novel genes support two models for gene birth: (1) *de novo* emergence ^54^ and (2) duplication degeneration ^55,56^. Determining a specific mechanism for gene birth for these auxiliary genes is beyond the scope of this work. However, reports of *de novo* gene origin in yeast and *Drosophila* show evidence of novel genes arising from previously short noncoding DNA sequences ^57,58^. Furthermore, small-scale duplications present an abundance of substrate for evolution to shape novel genes ^59^. One mechanism through which gene duplication leads to novel gene function is through neofunctionalization and/or subfunctionalization ^60,61^. After a gene duplicates, one copy can explore the evolutionary landscape without detrimental selective impacts because the other copy can compensate for loss or change of function. This can lead to fractionation of either or both copies and possibly reflect the shorter distribution of auxiliary genes.

We found that core and auxiliary enriched gene functions in *Vaccinium* align with observations in previous pangenome studies. Auxiliary genes are enriched for adaptive functions (eg defense response) similar to those of *Brachypodium distachyon*^22^ while core genes are enriched for basic cellular processes similar to those in *Brassica oleracea*^28^. Auxiliary genes are therefore targets of phenotypic differences between individuals and are of high agronomic importance. We discovered a large proportion of auxiliary genes in blueberry and cranberry that range between cultivar- to clade- to species-specific. This represents a substantial gene pool to leverage for future breeding efforts. Furthermore, genome-wide association studies (GWAS) can leverage pangenomes to discover genetic variants associated with important target traits ^34,36^. This may be a critical next step to further dissect the underlying genetics encoding important traits for *Vaccinium* breeding ^13^.

Lastly, the positionally-conserved core genes identified in this study can serve as foundational markers for the development of a genotyping platform for future molecular breeding efforts that is effective across a diversity of genetic backgrounds. In addition, certain auxiliary and/or species-specific genes, including those that contribute to improved disease resistance, stress tolerances and fruit quality traits, may serve useful to include on a genotyping platform for the blueberry and cranberry breeding community. Future research efforts will be focused on further characterizing these auxiliary genes and identifying those that greatly impact important target traits in both blueberry and cranberry.

## Materials and Methods

### Genome sequencing, assembly, and annotation

For genomic sequencing, leaf tissue was collected from each of the cultivars selected to best represent the genetic diversity of cranberry, southern highbush blueberry and northern highbush blueberry. DNA was extracted using a DNeasy extraction kit (Qiagen, Hilden, Germany). DNA quantity was assessed using a Qubit (ThermoFisher, Waltham, Massachusetts). Genomic libraries were prepared with the HyperPrep Library construction kit from Kapa Biosystems (Roche, Basel, Switzerland). Libraries were sequenced on a NovaSeq 6000 (Illumina, San Diego, California) in the Michigan State University Research Technology Support Facility (MSU RTSF) using a NovaSeq S4 reagent kit for 151 cycles from each end to generate paired 150 nucleotide long reads.

Genomic reads were quality and adapter trimmed using trimmomatic version 0.38 ^62^. Reads were then used to generate a hybrid *de novo* and reference based genome assembly. This assembly method was described in detail previously including tool versions and command line options ^63^. Briefly, genomic reads were mapped to the reference genome generated previously for cranberry ^38^ and blueberry ^37^. Mapped reads were used to generate a consensus genome sequence iteratively for three rounds. Then, unmapped reads were collected and *de novo* assembled into synthetic long-reads. These long reads were combined with the consensus sequence and incorporated into the final genome assembly for each cultivar.

Several tissues were collected for RNA sequencing analysis, including young leaf, mature leaf, green berry, and mature berry. Total RNA was isolated using the RNAeasy extraction kit (Qiagen, Hilden, Germany). RNA quantity was assessed using a Qubit (ThermoFisher, Waltham, Massachusetts). RNA libraries were prepared according to the mRNA HyperPrep kit protocol (Roche, Basel, Switzerland). All samples were sequenced in the MSU RTSF Genomics core with paired-end 150 bp reads on an HiSeq 6000 system (Illumina, San Diego, CA). Reads were quality and adapter trimmed using trimmomatic version 0.38 ^62^. They were mapped to their respective genome assemblies using hisat2 version 2.1.0 ^64^. The resulting SAM files were sorted and converted to BAM files using PicardTools version 2.18.1 SortSam function. From these alignments, transcriptome assemblies were generated using Stringtie version 2.1.3 ^65^. These transcriptome assemblies were used later to guide the gene annotation.

Each genome was annotated for protein coding genes using the MAKER2 pipeline and several lines of evidence ^66^. Proteins from Araport11 ^67^ and transcriptomes generated above were used as evidence. We also included the *Vaccinium corymbosum* ‘Draper’ CDS predictions ^37^ as evidence. We generated two *ab initio* models trained on ‘Draper’ gene models, SNAP and Augustus. Augustus models were generated using the script ‘train_augustus_draper.sh’ on a subset of 4,000 randomly selected gene models. SNAP models were generated using the script ‘train_snap_draper.sh’.

### Transposable element annotation

EDTA v2.0.0 was used to generate a pan-genome TE annotation ^68–76^. Default parameters were used in all cases except for the usage of the ‘--sensitive 1’ parameter which employs RepeatModeler to identify remaining TEs. First, individual repeat libraries were generated independently for each genome. Then, these libraries were filtered and combined using EDTA’s ‘make_panTElib.pl’ script to generate a pangenome repeat library for each genome group. Finally, the pangenome repeat library was used to re-annotate each source genome. Scripts and documentation for this analysis can be found at: https://github.com/sjteresi/Vaccinium_Pangenome_TE_Analysis.

### Identification of core and auxiliary genes

We aligned each genome using progressiveCactus to obtain whole genome alignments to identify positionally conserved core genes and identify missing auxiliary genes in cranberry ^43,44^. The progressiveCactus alignment also includes the highbush blueberry genomes. For blueberry, SynMap within CoGe using LAST and default parameters against the diploid W85 blueberry genome was used to identify positionally conserved core genes and auxiliary genes ^77^. We then identified orthologs between all cranberry and all blueberry proteomes using Orthofinder2 version 2.4.1 using default parameters (only the working directory specified) ^44^. As Orthofinder2 might identify members of the same gene family as orthologous, we decided to filter out any ortholog calls without an alignment within 5 kilobase-pairs of each other. We used the ‘filter_orthofinder2.sh’ script for ortholog calls.

### Gene statistic calculations

We calculated several feature values for each gene model including: gene length, coding sequence length, exon count, intron count, exon length, and intron length. These values were calculated using the ‘annotate_core_genes_vacc_pan.py’ script. Expression values were calculated using Kallisto v0.46.1 (Bray et al. 2016).

### Functional enrichments

Each proteome was functionally annotated using InterPro Scan version 5.28-67.0 ^78^. We converted the InterPro Scan annotation ID to a gene ontology ID using a manually curated translation table. We performed gene ontology term enrichment difference between core and auxiliary genes in R using the script ‘vacc_pan_go_enrichment.Rmd’ with the topGO package.

### Pangenome modeling

We modeled the core and auxiliary genomes of both blueberry and cranberry pangenomes based on the orthofinder results. We parsed the ‘Orthogroups.csv’ and ‘Orthogroups_UnassignedGenes.csv’ files using a custom script ‘model_pangenome_orthofinder.py’. This script calculates the number of core and auxiliary genes for each combination of individuals from 1 to the number of total accessions. We then plotted the average of core and auxiliary genes for each possible combination of accessions in Figure 2.

## Data availability

Custom scripts for analyses performed throughout this manuscript are available on Github (https://github.com/Aeyocca/VaccPan). Raw sequencing data are available on the NCBI SRA under project code PRJNA687008. Genome assemblies and annotations are available on the Genome Database for Vaccinium (GDV). Pangenome matrices are available on dryad XXXXXX.

## Supplementary information

### Figures

**Figure S1:**
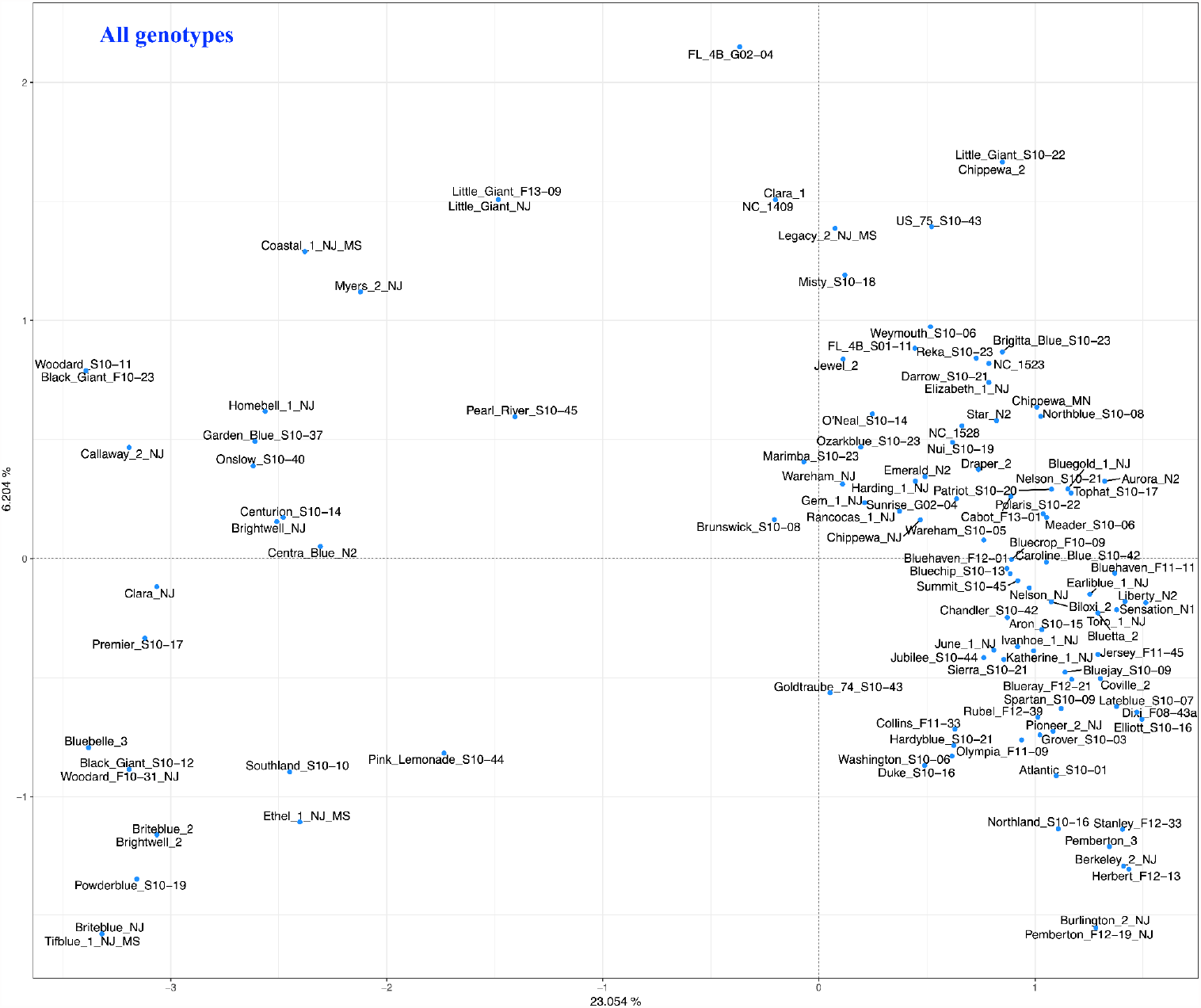
Principal component analysis of blueberry genotypes based on simple sequence repeat (SSR) markers.

### Tables

**Table S1:** Genome assembly statistics for blueberry and cranberry individuals assembled in this manuscript. This table is provided as a separate file.

**Table S2:** Gene ontology enrichment results obtained using Fisher’s exact test within the topGO R package. This table is provided as a separate file.

**Table S3:** Summary of structural variants in the ProgressiveCactus genome alignment. This table is provided as a separate file.

## Acknowledgements

This work was supported by Michigan State University AgBioResearch, Michigan State University Institute for Cyber-Enabled Research, NIH #5T32GM110523-10, NSF NRT-HDR #1828149 USDA-NIFA HATCH MICL02742, USDA-NIFA AFRI #1015241, and USDA-NIFA SCRI Award #2019-51181-30015. This work is supported in part by the National Science Foundation Research Traineeship Program (DGE-1828149) to MJ.

